# A database of biological thermal performances

**DOI:** 10.1101/2025.10.24.684478

**Authors:** Amahury J. Lopez-Diaz, José Ignacio Arroyo, Christopher Kempes, Alejandro Maass, Carlos Gershenson, Pablo Marquet, Geoffrey West

**Affiliations:** School of Systems Science and Industrial Engineering, Binghamton University, 4400 Vestal Pkwy E, Binghamton, NY 13902, USA; Santa Fe Institute, Santa Fe NM, USA; Center for Mathematical Modeling, University of Chile and IRL-CNRS 2807 Santiago, Chile; Millennium Institute Center for Genome Regulation, Santiago, Chile; Department of Mathematical Engineering, University of Chile, Santiago, Chile; Pontificia Universidad Católica de Chile

## Abstract

We built a database of thermal performance curves that includes different variables —mostly population growth rates —for species in representative groups across major domains, including viruses, bacteria, archaea, unicellular eukaryotes (such as fungi), and multicellular eukaryotes (including plants and both invertebrates and vertebrates). To build the database, we integrated previous databases and compiled additional data from individual studies, all of which were normalized to the same units when corresponding. Our database contains data for 1300 curves across 1125 different species and 38 variable types. The database consists of four (.csv) tables and an R code that can be explored through an R package or a Shiny app. Applications of our dataset include providing a resource to studies in thermal ecology and integration with other types of data, such as geographic temperatures. This would allow us to make further predictions, such as potential species distributions based on their thermal tolerances and their changes with climate warming.

## Background and Summary

Among all environmental parameters that affect biological processes, temperature is a universal predictor of biological processes, from the enzymatic to the ecosystem level, given the role of temperature in the regulation of the catalysis of biochemical reactions [1–3]. Remarkably, the response of organisms to temperature is similar from the molecular to the ecosystem level. The characteristic temperature response of biological quantities is exponential up to a critical point after which quantities decrease, and describe a right-skewed, symmetric, or left-skewed shape, but despite this, it has been theoretically- and empirically shown that it is a universal behavior common at all levels of organization [1]. At the individual/organismal level, the response of temperature to various traits, such as mobility speed, respiration, and fecundity, is known as “thermal performance”. Some of the properties, often called thermal traits, of the thermal performance that have been historically characterized include the minimum temperature (*T*_*min*_), maximum temperatures *T*_*max*_ tolerated by organisms, and the optimum *T*_*opt*_, which is the temperature at which the response reaches the highest value. These three fundamental traits are often called the cardinal temperatures. From these three fundamental limits, some additional thermal traits can be defined, such as the thermal breadth, which is often defined as the range of tolerated temperatures when the individuals reach half of their maximum performance, or the range, which is the difference *T*_*ran*_ = *T*_*max*_ − *T*_*min*_. Additional terms have been defined, such as the left range (*T*_*lran*_), which is the difference between the optimum and minimum, and the right range(*T*_*rran*_), which is the difference between the maximum and optimum, previously called in some articles thermal safety margin [4].

Quantifying a species’ thermal performance has practical applications such as predicting the optimal geographic region of growth of economically important species, predicting responses to climate change, or simply predicting the potential geographic range of a species [5–8]. For instance, for agronomically important species such as crops, it would be informative to know a species’ temperature performance and integrate it with average temperatures across Earth regions, which could predict the regions where a given species will grow optimally.

A few recent efforts have been made to compile data on the temperature performance of different species [9–13], but there are still many limitations. For example, some databases contain only a few thermal traits, rather than the full data describing the thermal performance of an individual or species. Others include data of a limited set of taxonomic groups, or types of variables, e.g., only population growth rate. On the other hand, those that include different variables do not include information on the thermal traits.

To overcome these limitations here we: i) compile in a single database data of thermal performance curves for different individual variables related to performance, ii) include data for all taxonomic groups of the Tree of Life, from viruses to mammals iii) include not just the data but also the thermal traits, iv) we provide user interfaces in the form of a Shiny app, and v) we also provide an R package to access to the data in the R environment. Applications of such datasets can be varied. One is providing a resource for studies in thermal ecology. The data can also be integrated with other types, e.g., geographic temperatures, to make further predictions, such as the potential distribution of a species based on its thermal tolerances.

## Methods

To construct our database of thermal performance curves (TPCs), we systematically compiled data from three main existing datasets—[14], [15], and [16]—and complemented these with primary data extracted manually from the literature. Our main curation criterion was to retain only complete or nearly complete TPCs, defined as curves containing at least three distinct temperature-response observations. This choice ensures sufficient resolution for reliably estimating thermal traits and prevents biases introduced by truncated or partial data, such as purely exponential or decay phases. Our dataset distinguishes itself by (1) including whole curve profiles (not just the thermal traits like optimum temperature), and (2) covering a broad taxonomic range, from viruses and bacteria to mammals, and a functional range of organismal responses, including per capita growth rate and other 37 performance metrics.

We obtained the data directly from tables or supplementary materials, when available. When data were not available in tables in the main text or supplementary material, and were only available in figures, we used WebPlotDigitizer (https://automeris.io/WebPlotDigitizer/) to extract high-resolution temperature-response datapoint pairs. Each record was verified for plausibility and completeness. To enable cross-study comparisons, we homogenized the units for each response variable using https://www.unitconverters.net. For each variable type (e.g., intrinsic growth rate), we identified the most commonly reported unit (e.g., 1/min for growth rates) and converted all entries accordingly.

Inclusion criteria required a minimum of three non-missing data points per curve. We included both concave and non-concave (i.e., monotonically increasing) curves but excluded fragments lacking a discernible peak or decay phase. Additionally, we discarded datapoints that fell outside the interquartile range (IQR) to reduce the influence of outliers. To define the minimum and maximum thermal traits (*T*_*min*_, *T*_*max*_), we used the lowest and highest temperature values for which the associated rates were within 5% of the smallest and largest performance variable values, respectively. All trait computations were implemented in R using custom scripts.

Species-level identifiers (IDs) were constructed using a combination of binomial names and, when needed, the first author’s last name and year of publication to disambiguate repetitions. Repeated specimens are limited to bacterial taxa, for which the literature provides abundant, heterogeneous data. We retained all variations (1300 records total) to support future comparative analyses and eventually plan to develop selection criteria to identify a “best representative” curve per species. The total number of unique species covered is 1125.

The database is stored in a structured relational format across four tables (Excel sheets):

- taxonomy: Contains columns id, domain, kingdom, phylum, binomial, name, and reference (DOI or citation string).
- temperature: Contains species IDs and associated temperatures (all the data is reported in ^*?*^C).
- response: Contains IDs, variable type, unit, concavity classification, and the response values aligned to temperature.
- traits: Contains derived thermal traits, including *T*_*opt*_ (optimum), *T*_*min*_, *T*_*max*_, *T*_*ran*_, *T*_*lran*_, and *T*_*rran*_.

This relational structure ensures internal consistency, facilitates SQL integration, and supports programmatic access through our provided R interface.

## Data Records

The database was built manually according to a relational model (Figure 1, Table 1). It has four tables. The first one contains everything related to taxonomy. Every organism has an associated ID, generated from the species’ binomial name. In the case of repeated specimens, we also add both the last name of the first author of the article from which the curve was extracted and the year of publication, so that the identifier can distinguish repetitions. Of course, there can be different response curves for a given species, and to distinguish them, we added an extra integer to the ID. In that first table, we also list basic taxonomic information such as domain, kingdom, phylum, and (binomial) scientific name. In addition, we added the reference from which the response curve for each organism was obtained. This reference is given in terms of a DOI, but it can also be a string with the citation in APA.

**Table 1.** Description of the variables in the tables of the dataset.

| Name | Description |
| --- | --- |
| Taxonomy |  |
| Domain | Taxonomic Domain; Bacteria, Archaea, Eukarya |
| Kingdom | Kingdoms; Bacteria, Archaea, Protozoa, Chromista, Fungi, Animalia, Plantae |
| Phylum | Taxonomic Phyla, 56 different Phyla |
| Scientific Name | Binomial taxonomic name |
| ID | A unique identifying string for the curve |
| Reference | Bibliographic reference of the source of the data |
| Temperature |  |
| ID | A unique identifying string for the curve |
| Temperature | Numeric data containing the temperature data |
| Response |  |
| ID | A unique identifying string for the curve |
| Response value | Numeric data of the performance variable |
| Variable name | Name of the performance variable, e.g., population growth rate. |
| Units | Units of the performance variable |
| Type | Concave or Not Concave curve |
| Thermal traits |  |
| ID | A unique identifying string for the curve |
| Optimum | Optimum temperature of the response curve |
| Minimum | Minimum tolerance temperature of the response curve |
| Maximum | Maximum tolerance temperature of the response curve |
| Range | Difference between maximum and minimum temperatures of the response curve |
| Left range | Difference between optimum and minimum temperatures of the response curve |
| Right range | Difference of maximum and optimum temperatures of the response curve |

**Figure 1.**
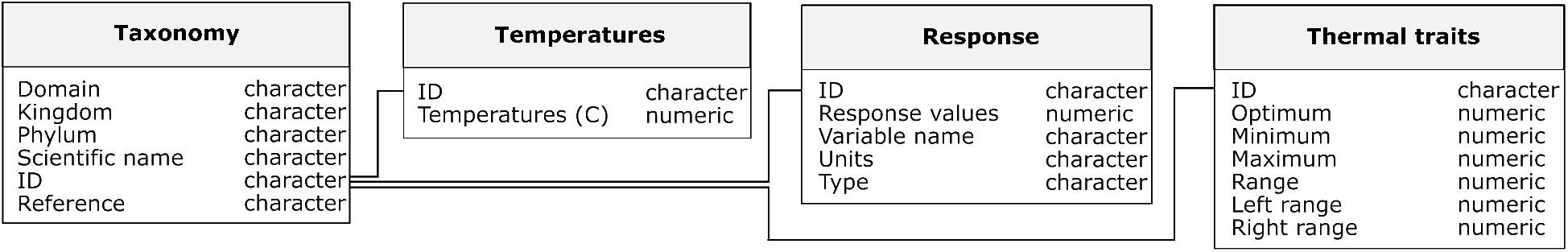
Database schema depicting the four tables that compose our dataset. See Table 1 for a description of each of the variables.

The second table contains the respective temperatures of the performance curves. All this data is given in Celsius degrees. We also included the ID to indicate which organism each temperature belongs to.

The third table contains the corresponding performance values for every temperature shown in the second layer. In addition to the ID, we included the variable type (to indicate which type of response we are referring to), the unit associated with this variable type, and the concavity, this latter being “Concave” if the curve is concave (i.e., has a well-defined optimum) or “No Concave” if the curve is monotonically increasing.

The fourth table summarizes the thermal traits associated with the response curves. We again indicate the ID and, depending on the concavity, display the minimum, maximum, optimum, and distinct thermal ranges, as defined in the introduction.

The total number of specimens in the database is 1300, counting repeated organisms for a given performance trait. All the repeated species are bacteria, and the variations among the curves are shown in the supplementary material. The total number of repeated species is 175, with *Escherichia coli* being the most frequent with 22 repetitions. Counting the non-repeating organisms for each performance trait yields a total of 1125 species. The taxonomic distribution and allocation of responses in our database are summarized in Table S2. The total number of performance traits that our database compiles is 38. Of these, three stand out: intrinsic growth rate, photosynthetic rate, and running speed. For these traits, there are a considerable number of curves, totaling 1055 for growth rate, 114 for photosynthetic rate, and 51 for running speed, as indicated in Table S2.

## Technical Validation

Each curve underwent both manual and programmatic validation steps to ensure internal consistency and biological plausibility. Thermal trait values (*T*_*min*_, *T*_*opt*_, *T*_*max*_) were cross-checked using the constraint: *T*_*min*_ *< T*_*opt*_ *< T*_*max*_. Any violations triggered a manual re-evaluation of the curve or its classification.

To validate the concavity designation, we used the function get curves(), which compares the temperature and response sheets. Curves were labeled as *Concave* if at least one temperature yielded a response lower than the peak, indicating a decline after the optimum. Monotonically increasing curves (typically microbial growth rates under suboptimal conditions) were labeled as *No Concave*.

Outlier filtering was performed by computing the interquartile range of the response values and removing datapoints lying beyond 1.5×IQR from the first and third quartiles. This ensured that extreme fluctuations due to measurement error or digitization artifacts did not bias trait extraction.

Finally, to further corroborate that the critical temperature limits were consistent, we evaluated distributions and correlations among the so-called cardinal temperatures: the *T*_*min*_, *T*_*opt*_, and *T*_*max*_. The distributions of the cardinal temperatures were all within the well-known temperature range of life: ≈ -20 to 120 degrees Celsius [14], and exhibited a bimodal distribution, most likely defining meso- and thermophiles. It has been reported in the literature that these limits are correlated among them (e.g., [17]. The (Pearson) correlations (coefficients) among these three thermal traits were high and significant in all cases.

All dataset entries are traceable via the reference column in the taxonomy sheet, which lists either a DOI or citation string for every species-curve. These references ensure full transparency and reproducibility. We are currently working to formalize version control with Git and plan to offer updated releases through a persistent digital repository.

## Usage Notes

Users should be aware that, although each curve represents a well-defined thermal response, intrinsic variability arises from differences in experimental protocols, medium composition, and strain-level variation (especially among microbes). In particular, we retained multiple curves for frequently studied bacterial species such as Escherichia coli (22 distinct records), where curve shapes and optima can differ substantially across studies. In future iterations, we plan to derive consensus curves via data quality scoring, weighted averaging, or normalization. This version of the database is strongly dominated by growth rate responses (≈ 81%), with secondary representation for photosynthetic rates, running speed, filtration rate, and oxygen consumption (Table S1). Future expansions will prioritize the inclusion of underrepresented trait types, such as metabolic or parasitism rates, which are key for ecological forecasting. While the dataset already spans a wide array of taxa, some clades — particularly those of agricultural, ecological, or industrial importance — are underrepresented (Table S1). We aim to expand coverage for such groups in upcoming versions, with attention to species relevant to food production, carbon cycling, or conservation. The database supports numerous applications across ecology, evolutionary biology, and applied systems biology, including i) Predicting responses to climate warming —changes in abundance and distribution, range shifts, or species persistence — using physiological tolerance limits; calibrating metabolic theory models; and screening temperature optima for industrial strains in biotechnology and agriculture. We developed the R package “TBio”, which includes the data and a set of functions for exploring it. For now, the package is available upon request (and after publication, it will be available to be downloaded from GitHub). After downloading it in a local folder, the R package can be simply installed in the R environment using devtools::install(“TBio”) and uploaded using library(TBio). We also developed an interactive Shiny app user interface, “TBio Explorer”, which facilitates subsetting, plotting, and exporting curated performance curves. The modular database structure also supports integration with trait-based ecological models, phylogenetic frameworks, and global environmental datasets. The code to run the app locally is available upon request (and, after publication, will be available for download from GitHub). After downloading it in a local folder, the app can be simply run by following the instructions in the README file.

## Acknowledgements

NSF Award “Building and Modeling Synthetic Bacterial Cells” (Award Number 1840301), NSF Award “Towards a unified theory of regulatory functions and networks across biological and social systems” (Award Number 2133863), and Center for Mathematical Modeling (CMM), Grant FB210005, BASAL funds for Centers of Excellence from ANID-Chile.

## Author Contributions

ALD and JIA made the database and technical validation. JIA, ALD, CK, AM, CG, PM, and GW wrote the paper.

## References

1. Arroyo, J. I., Díez, B., Kempes, C. P., West, G. B. & Marquet, P.A. A general theory for temperature dependence in biology. Proceedings of the National Academy of Sciences 119, e2119872119 (2022).

2. Dell, A. I., Pawar, S. & Savage, V. M. Systematic variation in the temperature dependence of physiological and ecological traits. Proceedings of the National Academy of Sciences 108, 10591–10596 (2011).

3. Brown, J. H., Gillooly, J. F., Allen, A. P., Savage, V. M. & West, G. B. Toward a metabolic theory of ecology. Ecology 85, 1771–1789 (2004).

4. Sunday, J. M. et al. Thermal-safety margins and the necessity of thermoregulatory behavior across latitude and elevation. Proceedings of the National Academy of Sciences 111, 5610–5615 (2014).

5. Sinclair, B. J. et al. Can we predict ectotherm responses to climate change using thermal performance curves and body temperatures? Ecology letters 19, 1372–1385 (2016).

6. Khelifa, R., Blanckenhorn, W. U., Roy, J., Rohner, P. T. & Mahdjoub, H. Usefulness and limitations of thermal performance curves in predicting ectotherm development under climatic variability. Journal of Animal Ecology 88, 1901–1912 (2019).

7. Kingsolver, J. G. & Buckley, L. B. Quantifying thermal extremes and biological variation to predict evolutionary responses to changing climate. Philosophical Transactions of the Royal Society B: Biological Sciences 372, 20160147 (2017).

8. Wagner, T. et al. Predicting climate change impacts on poikilotherms using physiologically guided species abundance models. Proceedings of the National Academy of Sciences 120, e2214199120 (2023).

9. Diamond, S. E., da Silva, C. R. & Medina-Báez, O. A. A multicontinental dataset of butterfly thermal physiological traits. Scientific Data 11, 1348 (2024).

10. Valente, S. & Colloca, F. A dataset of thermal preferences for Mediterranean demersal and benthic macrofauna. Scientific Data 11, 314 (2024).

11. Pottier, P. et al. A comprehensive database of amphibian heat tolerance. Scientific Data 9, 600 (2022).

12. DuBose, T. P. et al. Thermal traits of Anurans database for the Southeastern United States (TRAD): a database of thermal trait values for 40 Anuran species. Ichthyology & Herpetology 112, 21–30 (2024).

13. Bennett, J. M. et al. GlobTherm, a global database on thermal tolerances for aquatic and terrestrial organisms. Scientific Data 5, 1–7 (2018).

14. Corkrey, R. et al. The biokinetic spectrum for temperature. PLoS One 11, e0153343 (2016).

15. Rezende, E. L. & Bozinovic, F. Thermal performance across levels of biological organization. Philosophical Transactions of the Royal Society B 374, 20180549 (2019).

16. Kontopoulos, D.-G., Sentis, A., Daufresne, M., Dell, A. I. & Pawar, S. No model to rule them all: a systematic comparison of 83 thermal performance curve models across traits and taxonomic groups. bioRxiv, 2023–09 (2023).

17. Rosso, L., Lobry, J. R. & Flandrois, J.-P. An unexpected correlation between cardinal temperatures of microbial growth highlighted by a new model. Journal of theoretical biology 162, 447–463 (1993).

