## Supplementary Material for "A database of biological thermal performances"

### Supplementary Material for
### A database for biological thermal performances

Amahury J. Lopez-Diaz† (1), José Ignacio Arroyo† (2), Christopher Kempes (2), Alejandro Maass (3,4,5), Carlos Gershenson (1), Pablo Marquet (2, 3, 6), and Geoffrey West (2)

(1) School of Systems Science and Industrial Engineering, Binghamton University, 4400 Vestal Pkwy E, Binghamton, NY 13902, USA
(2) Santa Fe Institute, Santa Fe NM, USA.
(3) Center for Mathematical Modeling, University of Chile and IRL-CNRS 2807 Santiago, Chile.
(4) Millennium Institute Center for Genome Regulation, Santiago, Chile.
(5) Department of Mathematical Engineering, University of Chile, Santiago, Chile.
(6) Pontificia Universidad Católica de Chile

† These authors contributed equally to this work

Corresponding Author(s):
J. Ignacio Arroyo:,
Amahury Lopez:

† These authors contributed equally to this work

Corresponding Author(s):
J. Ignacio Arroyo:,
Amahury Lopez:

Table S1. Description of the variables in the tables of the dataset.

| | | | Number of Curves | Percent |
|---|---|---|---|---|
| | | Domain | Kingdom | |
| Virus | | | 15 | 1.1538% |
| | Bacteria | Bacteria | 527 | 40.5384% |
| | Archaea | Archaea | 172 | 13.2307% |
| | Eukarya | Fungi | 35 | 2.6923% |
| | Eukarya | Protozoa | 21 | 0.0161% |
| | Eukarya | Chromista | 66 | 0.0507% |
| | Eukarya | Plantae | 140 | 10.7692% |
| | Eukarya | Animalia | 324 | 24.9230% |
| | Response type | | | |
| | Population growth rate | | 1055 | 81.1538% |
| | Photosynthetic rate | | 114 | 8.7692% |
| | Running speed | | 51 | 3.9230% |
| | Filtration rate | | 16 | 1.2307% |
| | Oxygen consumption | | 11 | 0.8461% |
| | Other | | 53 | 4.0769% |
